# An engineered lactate responding promoter system operating in glucose-rich and anaerobic environments

**DOI:** 10.1101/2021.01.06.425364

**Authors:** Ana Zúñiga, Hung-Ju Chang, Elsa Fristot, Jerome Bonnet

## Abstract

Bacteria equipped with genetically-encoded lactate biosensors would support several applications in biopharmaceutical production, diagnosis, or therapeutics. However, many applications involve glucose-rich and anaerobic environments, in which current whole-cell lactate biosensors have low performance. Here we engineered a synthetic lactate biosensor system by repurposing the natural LldPRD promoter regulated by the LldR transcriptional regulator. We removed glucose catabolite repression by designing a hybrid promoter, containing LldR operators and tuned both regulator and reporter gene expression to optimize biosensor signal-to-noise ratio. The resulting lactate biosensor, termed ALPaGA (**<u>A</u>** <u>**L**</u>actate <u>**P**</u>romoter Oper<u>**a**</u>ting in <u>**G**</u>lucose and **A**naerobia) can operate in glucose rich, aerobic and anaerobic conditions. Our work provides a versatile lactate biosensing platform suitable for many environmental conditions.

## MAIN TEXT

Lactate is produced from anaerobic metabolism^1^ and has long been considered as a waste product. Lactate can negatively influence the production yield and quality of several bioprocesses and its monitoring is thus important in the food and biopharmaceutical industry^2–4^.

On the other hand, lactate is a versatile and important raw material for various industrial processes. Lactate derivatives are used as food additives for their antimicrobials, antioxidants, or flavoring properties^5^. Lactate is also a basic building block for various biopolymers^6–8^ such as polylactic acid used in the construction of biomedical devices because of its biodegradability and biocompatibility^9^. Lactate production is thus an important part of the bioeconomy and is mostly produced from renewable feedstocks using the natural sugar fermentation capacity of a wide number of microbes and fungi^10^.

As a central product of anaerobic metabolism, lactate is also a key biomarker of the human physiological state^1^. In medicine, lactic acidosis occurs in several conditions such as sepsis or diabetes and is an important parameter to be monitored in patients admitted in intensive care units^11^. In oncology, lactate produced by cancer cells is a hallmark of solid tumors that leads to tumor acidification and participates in immune system inhibition^12^.

For all these reasons, lactate monitoring is important and several detection systems have been developed^13–15^. Most of them involve enzymatic reactions of lactate oxidase and lactate dehydrogenase coupled to amperometric detection^16^ or electrochemical biohybrid oxygen sensor based on natural bacteria metabolism^17^. Yet, these biosensing methods either have low sensitivity or are expensive, limiting their use and deployment.

Another approach for lactate detection is to use whole-cell biosensors. These sensors based on living cells, often bacteria, generally use a specific transcription factor responding to a signal of interest and its target promoter to regulate the expression of a reporter gene^18,19^. This strategy has produced a wide range of biosensors responding to a variety of metabolites including heavy-metals, butanol, alkanes, acyl- or malonyl-CoA^20–27^. Whole-cell biosensors are highly sensitive, specific, and the replicating nature of microorganisms support their cost-effective production. In addition, genetically-encoded sensors can also serve as input signals for genetic circuits controlling cellular behavior such as cell growth in specific environmental conditions^28^, conditional control and optimization of metabolic pathways^29,30^, or production of a therapeutic payload^31, 32, 31^.

Genetically encoded lactate biosensors operating in bacteria have been recently engineered for monitoring lactate levels in biopharmaceutical production or restrain the growth and activity of bacterial cancer therapeutic to the tumor microenvironment^4,33,34^. All these biosensors are based on the *Escherichia coli* LldPRD promoter controlled by the LldR regulator in response to lactate^35,36^ (**Figure 1A**). LldR triggers induction of the lldPRD operon responsible for lactate metabolism when *E.coli* cells are grown in lactate as sole carbon source. Despite having demonstrated functionality and promising results, existing lactate biosensors face several challenges.

**Figure 1.**
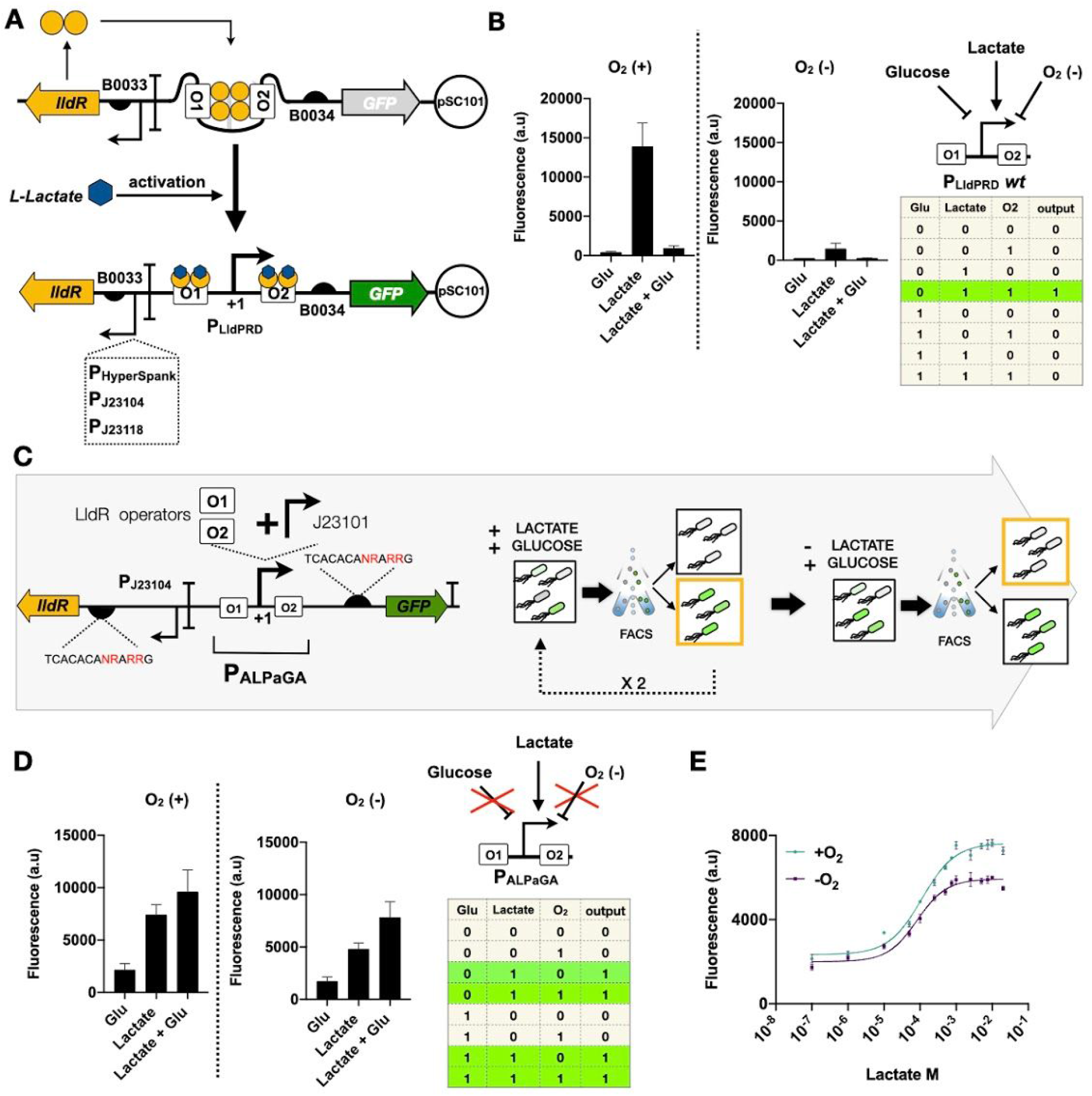
Engineering a L-lactate whole-cell biosensor operating in glucose-rich and anaerobic environment. **(A)** Architecture and regulation of the low-copy lactate responsive biosensor based on the wt LldPRD system. **(B)** Biosensor response to lactate, in presence or not of 0.4% glucose and oxygen. (left) response of wt LldPRD promoter to 0.4% glucose, 10 mM of lactate and both under aerobic (+O_2_) or anaerobic (−O_2_) conditions. (right) Regulatory logic diagram of the wild type P_LldPRD_ promoter response to lactate, glucose and oxygen. Truth table is represented below. **(C)** Design and optimization of **<u>A</u>** <u>**L**</u>actate <u>**P**</u>romoter Oper<u>**a**</u>ting in <u>**G**</u>lucose and **A**naerobia, P_ALPaGA_. *(left)* Design of the synthetic promoter, J23101 was used as a constitutive core promoter and the two operators were included conserving the original distance from the LldPRD promoter. The RBS library for lldR and GFP is detailed. *(right)* schematic representation of library screening by FACS enrichment in M9 plus glucose in presence and absence of lactate. **(D)** Engineered sensor response to combinations of lactate, glucose and oxygen. *(left)* response of synthetic promoter system to 0.4% glucose, 10 mM of lactate and its combination under aerobic (+O_2_) or anaerobic (−O_2_) conditions. *(right)* Regulatory logic diagram and truth table of the ALPaGA promoter system. **(E)** Dose response of the engineered lactate biosensor to lactate and glucose. The fluorescence (a.u.) is shown under aerobic (+O_2_) or anaerobic (−O_2_) conditions. For all experiments, the data represent the average of three biological replicates performed on different days in triplicates. Errors bars: +/− SD

First, current lactate biosensors operate on high-copy number plasmids, which are notoriously associated with metabolic burden^37,38^ and genetic instability^39^, limiting their application, both *in vitro^40^* and *in vivo^41^*. Biosensors operating at low-copy numbers are thus needed. Second, for many applications, the environment is rich in glucose, the preferred carbon source for *Escherichia coli^42^* which often shuts down operons controlling the utilization of other carbon sources though carbon catabolite repression (CCR)^43–46^. Indeed, the native lactate utilization operon is subject to CCR^35^ and at least one of the previously engineered lactate biosensors was shown to exhibit a lower performance and a ~70% lower induction response in presence of glucose^4^. Third, lactate biosensors would be highly useful in anaerobic environments to monitor lactate production. For example, the best production of lactate is obtained from anaerobically growing lactic acid bacteria^10^ and lactate production in solid tumors is linked to their hypoxic nature^12^. Yet, transcription of the lldDRPD operon was shown to be repressed under anaerobic reducing conditions^47–50^.

To extend the range of application of lactate biosensors, we thus aimed at engineering a sensor operating in glucose-rich and anaerobic environments. By analyzing the regulatory logic of the lactate biosensor system based on the native LldRPD promoter operating at low-copy numbers in *E. coli,* we observed strong repression by glucose and anaerobic conditions. We then engineered and finely tuned a synthetic L-lactate biosensor able to operate in presence of glucose and under both aerobic and anaerobic conditions.

We first assessed the functionality of the L-lactate whole-cell biosensor by constructing the biosensor described by Goers and coworkers^4^. This biosensor is based on the wild-type promoter of LldPRD operon and expresses the LldR regulator from the pHyperspank promoter. To address the issues associated with high-copy numbers, we placed this system on a low-copy plasmid with pSC101 origin of replication (5-10 copies)^51^. We designed two other versions of the biosensor in which we used two different strong constitutive promoters to control expression of the *LldR* gene (**Figure 1A**). To assess the sensitivity of the biosensors to glucose-mediated carbon catabolite repression, we tested their response in M9 with or without 0.4% glucose (22 mM). All biosensors were able to sense *L*-lactate in M9 when lactate was used as a sole carbon source, demonstrating that this system can operate at low-copy numbers (**Figure 1B, Supplementary Figure S1**). The versions in which *lldR* expression was driven by strong constitutive promoters (in particular J23104) had a much better response than the one in which pHyperspank was used, after 4 h of induction. Sensor exhibited a ~7 fold change in accordance with previously published results^33,34^ with a half maximal effective concentration (EC50) of ~1.6 mM. However, when glucose or glycerol were added as a carbon source the biosensor response considerably dropped, confirming strong catabolic repression of the LldPRD promoter by these sugars, with no detectable response in the presence of glucose (**Figure 1B, Supplementary Figure S2**). Catabolic repression directly affects the LldPRD promoter, as repression is observed even when the pHyperspank promoter (also known to be subject to CCR) was not used to control *lldR* expression. We then tested the sensor response in anaerobic conditions. As expected from literature, we observed no response from our L-lactate biosensor after 16h of induction, confirming strong inhibition of the promoter (**Figure 1B, Supplementary Figure S2**). These results demonstrate that while capable of operating at low-copy numbers, the lactate biosensor based on the wild-type LldPRD system is not usable in glucose-rich nor in anaerobic conditions, greatly limiting its range of applications.

To overcome the catabolic repression observed by glucose, we engineered a synthetic L-lactate promoter. This promoter was constructed by using a sequence from a constitutive promoter to replace the sequence between the −35 and −10 of the wild type LldPRD promoter, combined with the operator sequences recognized by LldR. The first version of the system using this synthetic promoter was highly leaky (**Supplementary Figure S3**). To optimize the biosensor response through directed evolution we created a double RBS library to concomitantly variate expression of GFP and *lldR* (**Figure 1C**, left). The double library was transformed into *E. coli* DH5alpha and FACS sorting was used to screen variants based on GFP fluorescence intensity (**Fig. 1C**, right, **Supplementary Figure S4**). For the first sort, cells producing GFP were selected after induction with 20 mM of lactate *in presence of glucose* after overnight growth. A second round of positive selection was made by using 10 mM of lactate. A third and last round in the absence of inducer was performed to select variants with lower leakiness. After these sequential rounds, 80 biosensor variants were recovered and tested for their response to 1 mM of lactate and 0.4% glucose in aerobic and anaerobic conditions (**Figure 1D**, **Supplementary Figure S4**). Variants with the higher fold changes were selected and characterized as a function of L-lactate concentration. The final L-lactate sensor variant had a ~3.4 fold change in the presence of glucose under aerobic conditions and a ~3.2 fold change in the presence of glucose under anaerobic conditions (**Figure 1D**).

We then established the dose-response curve of the biosensor to *L*-lactate in the presence of glucose under aerobic and anaerobic conditions (**Figure 1E**) and calculated an EC50 of ~110*μ*M under aerobic conditions and ~90*μ*M under anaerobic conditions. Quite surprisingly, when we tried various concentrations of glucose, we observed an increase in basal GFP fluorescence at higher glucose concentration (**Supplementary Figure S5**). We attribute this effect on the positive effect of higher glucose concentration on bacterial growth and metabolism (**Supplementary Figure S6**). Nevertheless this effect was small compared with the increment due to the lactate induction. We termed our promoter ALPaGA for “**<u>A</u>** <u>**L**</u>actate <u>**P**</u>romoter Oper<u>**a**</u>ting in <u>**G**</u>lucose and **A**naerobia”.

In conclusion, we engineered a synthetic lactate biosensor driven by the engineered ALPaGA promoter reliably operating in glucose-rich and anaerobic conditions in which previous systems using the wild-type LldPRD promoter had poor performance. In addition, we show that the biosensor can operate at a low-copy number, reducing potential metabolic burden effects, and making it compatible with future clinical applications. Our system still exhibits some background expression due to leakiness of the engineered promoter, negatively affecting its signal-to-noise ratio. While we were able to reduce leakiness *via* directed evolution of ribosome binding sites controlling regulator and reporter expressions, further improvement in biosensor signal-to-noise ratio could be done using other circuit engineering methods which have already been applied to the *wt* LldR system^34,52^.

The ALPaGA lactate biosensor presented here will be useful for many applications in which the environment is glucose rich and/or anaerobic, such as monitoring bioproduction processes or restricting the activity of bacterial cancer therapeutics within the tumor microenvironment.

## METHODS

### Strains and plasmids

The implementation of the biosensor was done in *E. coli* strain DH5alphaZ1 (laci^q^, PN25-tetR, Sp^R^, deoR, supE44, Delta(lacZYA-argFV169), Phi80 lacZDeltaM15, hsdR17(rK− mK+), recA1, endA1, gyrA96, thi-1, relA1). DH5alphaZ1 was grown on LB media with kanamycin 25μg/mL to do the cloning of plasmids. For experimental measurements the cells were grown in M9 minimal medium supplemented with 0.4% of glycerol and kanamycin 25μg/mL. L-lactic acid (Sigma-Aldrich, L1750) was used to induce the cells at different concentrations. Carbon sources concentration as glycerol and glucose, are given as % mass (w/v %).

### Library design, and plasmids construction

The RBS library design refers to iGEM parts: BBa K1725301 - K1725332 (Group: iGEM15_Glasgow). All variants are derived from Anderson-family RBS BBa_B0032 with the following sequences: TCACACANRARRG. One-step isothermal Gibson assembly was used to build all plasmids described.

All the biosensor parts were built on the backbone pSB4K5^51^, containing a pSC101 origin of replication and kanamycin resistance. Enzymes for the one-step isothermal assembly were purchased from New England BioLabs (NEB, Ipswich, MA, USA). PCR were performed using Q5 PCR master mix and One-Taq quick load master mix for colony PCR (NEB), primers were purchased from IDT (Louvain, Belgium), and DNA fragments from Twist Bioscience. All plasmids were purified using QIAprep spin Miniprep kit (Qiagen) and sequence verified by Sanger sequencing in Eurofins Genomics, EU.

DNA encoding the LldR transcription factor (lldR) and the wild type promoter sequence pLldPRD were amplified from *E. coli* based on ^4^ design. All primers were designed to support cloning by Gibson assembly at an identical location in pSB4K5 template vector. Consequently, all primers were composed of the 40 bp spacer 0 at 5′ end, and 40 bp spacer N at 3′ end. The DNA sequence for the Alpaga promoter was synthesized as linear fragments by Twist Bioscience. Each DNA fragment was PCR amplified and assembled between spacer 0 and N in pSB4K5 template vector. All DNA sequences are listed in Table for Supplementary Information.

### Sensor characterization

The different biosensor circuits were transformed in *E. coli* strain DH5alphaZ1 and plated on LB agar medium containing kanamycin. Three different colonies for each circuit were picked and inoculated, separately, into 500μL of M9 supplemented with 0.4% of glycerol and kanamycin in 96 DeepWell polystyrene plates (Thermo Fisher Scientific, 278606) sealed with AeraSeal film (Sigma-Aldrich, A9224-50EA) and incubated at 37°C for 16h with shaking (300 rpm) and 80% of humidity in a Kuhner LT-X (Lab-Therm) incubator shaker. After overnight growth the cells were adjusted at OD 0.1 in a fresh medium with antibiotics and L-lactate at different concentrations, with or without glucose 0.4%. Cells were induced at 37°C for 16h with or without shaking for aerobic and anaerobic conditions, respectively. The induction in anaerobic conditions was done by growing the cells in a BD GasPak EZ Anaerobe Container System (BD; 260003) with a BD GasPak EZ pouch system (BD; 260678) for 16h at 37°C, and analyzed by flow cytometry. All experiments were performed at least 3 times in triplicate.

The goodness of fit and the sensitivity were calculated by applying non-linear regression using sigmoidal curve function and EC50 using GraphPad Prism. The fold change was calculated as follows: for lactate only conditions, because cells needed lactate as a carbon source, we did not have a data point without lactate. Because the promoters are leaky, we could not either use negative control cells without biosensor as our non-induced condition. Thus, fold change was calculated as the fluorescence intensity at maximal lactate concentration divided by the fluorescence intensity at the lowest lactate concentration, which is well below the threshold lactate concentration at which an inflexion is observed in dose-response curves. For cells growing in glucose, fold change was calculated as fluorescence intensity at maximal lactate concentration divided by the fluorescence intensity when no lactate was added.

### Flow cytometry

Flow cytometry was performed on Attune NxT flow cytometer (Thermo Fisher) equipped with an autosampler and Attune NxT Version 2.7 Software. Experiments on Attune NxT were performed in 96-well plates with setting; FSC: 200 V, SSC: 380 V, green intensity BL1: 460 V (488 nm laser and a 510/10 nm filter). All events were collected with a cutoff of 20,000 events. Every experiment included a negative control harboring the plasmid but without reporter gene, to generate the gates. The cells were gated based on forward and side scatter graphs and events on single-cell gates were selected and analyzed, to remove debris from the analysis (Fig. S21), by Flow-Jo (Treestar, Inc) software. The geometrical median of the fluorescence histogram of each gated population was calculated and is reported here as the fluorescence value of a sample in arbitrary units (a.u.).

### Cell sorting

Cell sorting was performed using a Bio-Rad S3 cell sorter (Bio-rad). 100,000 cells were gated under three different induction conditions (Fig. S5). They were collected in SOC medium during the sorting and recovered for 1 hour before being inoculated in 10 mL of LB/chloramphenicol medium for 18 hours at 37°C with shaking.

## Supporting information

DNA sequences

## ACKNOWLEDGMENTS

We thank members of the synthetic biology group and of the CBS for fruitful discussions and feedback. This work was supported by an ERC starting grant “COMPUCELL” and ANR “SynbioDiag”. J.B. also acknowledges the INSERM Atip-Avenir program and the Bettencourt-Schueller Foundation for continuous support. The CBS acknowledges support from the French Infrastructure for Integrated Structural Biology (FRISBI) ANR-10-INSB-05-01.

## AUTHORS CONTRIBUTION

A.Z. and J.B. designed the research. A.Z. and H.J.C. designed variants libraries and performed sorting procedure for biosensor optimization by directed-evolution. A.Z. and E.F. designed and performed experiments in anaerobic conditions. All other experiments were performed by A.Z. A.Z. and J.B. analyzed the data and wrote the article. All authors reviewed and approved the manuscript.

