## Supplementary Materials for "An engineered lactate responding promoter system operating in glucose-rich and anaerobic environments"

**Supplementary Figures**

Supplementary Figure 1: Characterization of the different versions of L-lactate biosensor based on wild-type LldPRD system in aerobic conditions

Supplementary Figure 2: Effect of glucose and anaerobic conditions on the performance of the L-lactate biosensor based on wild-type LldPRD.

Supplementary Figure 3: Characterization of the first version of the synthetic L-lactate promoter showing high-leakiness

Supplementary Figure 4: Schematics of FACS enrichment and screening of the RBS library.

Supplementary Figure 5: Two-dimensional response of the ALPaGA L-lactate biosensor to increasing concentrations of lactate and glucose in aerobic (+$O_2$) and (-$O_2$) anaerobic conditions.

Supplementary Figure 6: Effect of glucose on the growth of the ALPaGA L-lactate biosensor

**Supplementary Tables**

Supplementary table 1: Plasmid parts list and Plasmid sequences

**Figure S1**

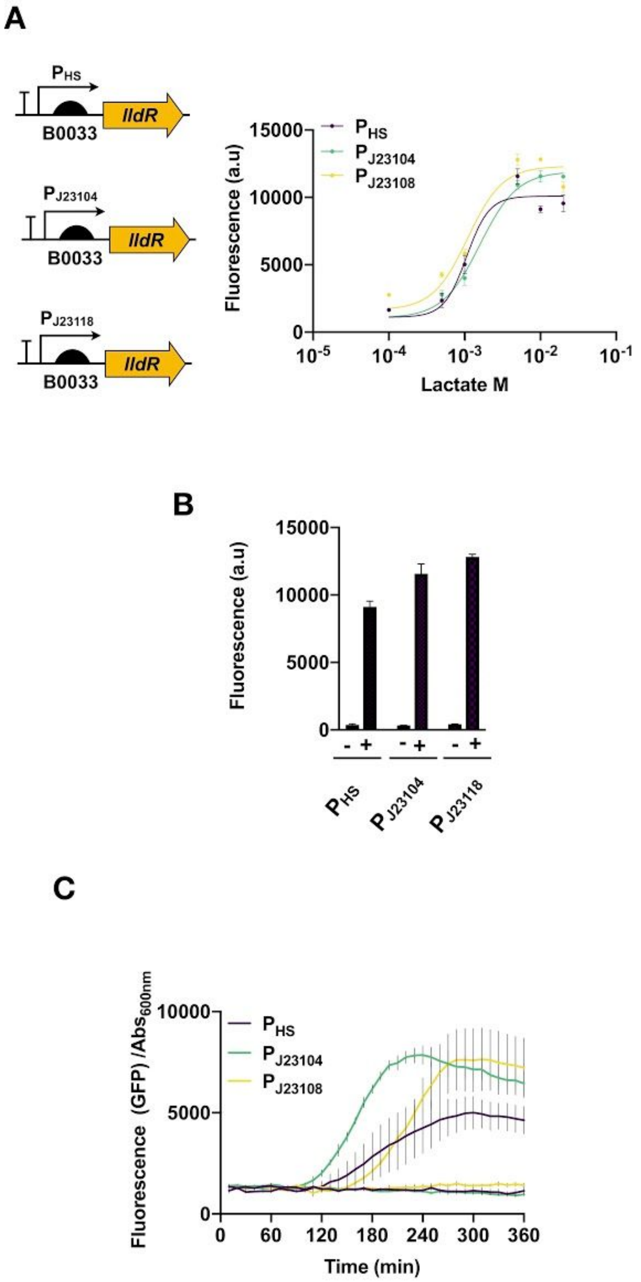


**Figure S1. Characterization of the different versions of L-lactate biosensor based on wild-type LldPRD system in aerobic conditions (A)** *(left)* Diagrams of the different constructions of the biosensor by using different constitutive promoters for lldR expression; *(right)* response function for each biosensor. Error bars represent the standard deviation to the mean of three biological replicates performed in triplicates on different days. **(B)** Fluorescence in arbitrary units and biosensors in response to 10 mM of lactate in aerobic conditions. The graphs represent the mean of the fold change of three replicates in the presence or not of lactate. **(C)** Dynamic of inductions represented by the fluorescence normalized by absorbance (OD600nm). Cells were induced with 10mM of lactate for 16 hours. Error bars represent the standard deviation of three biological replicates performed in triplicates on different days. HS; hyperspank.

**Figure S2**

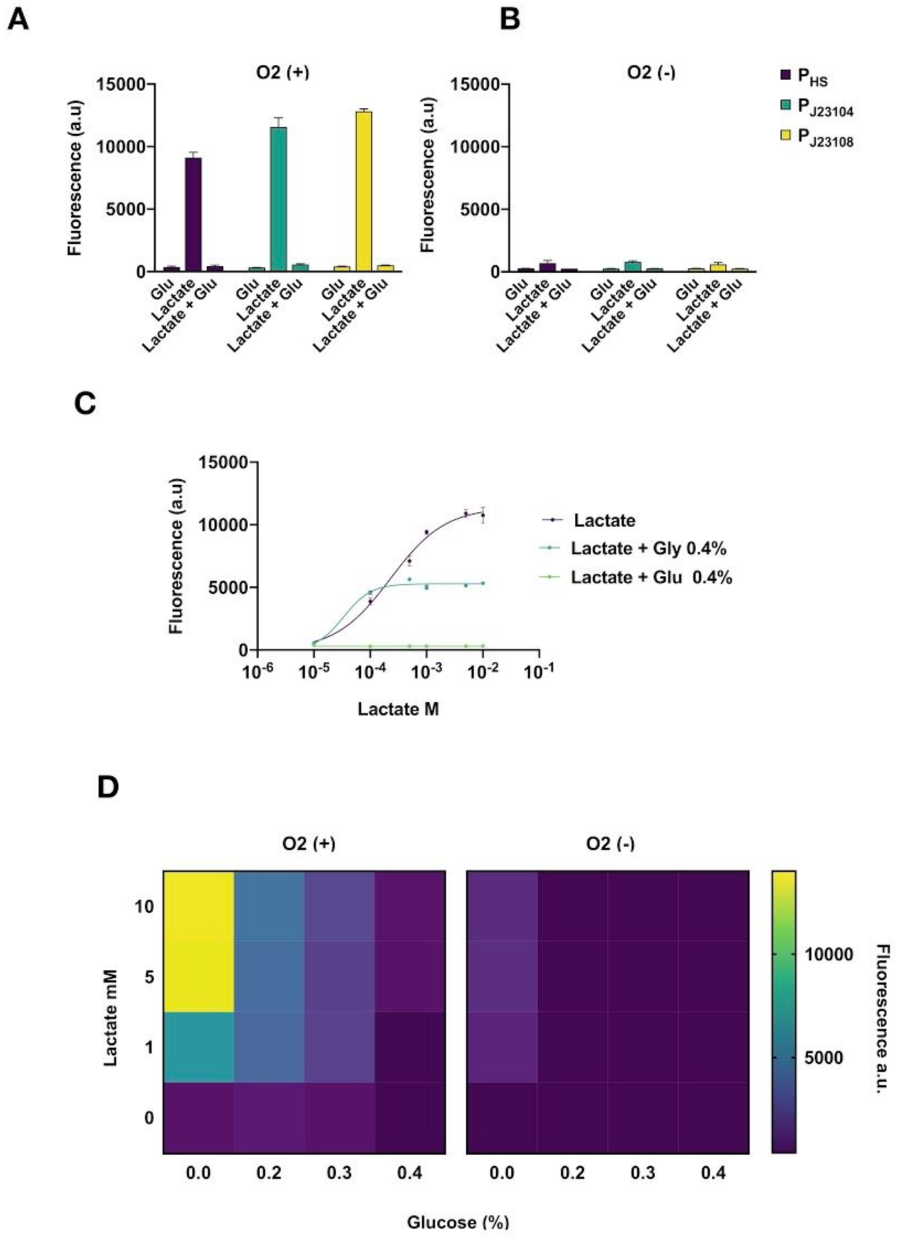


**Figure S2. Effect of glucose and anaerobic conditions on the performance of the L-lactate biosensor based on wild-type LldPRD.** Fluorescence in arbitrary units (a.u.) of biosensors in response to 10 mM, in presence of 0.4% of glucose, under aerobic **(A)** or anaerobic **(B)** conditions. The graphs represent the mean of the fold change of three replicates. **(C)** Effect of 0.4% glucose and 0.4% glycerol on the response of PJ23104-LldR biosensors to L-lactate after 4 hours. Error bars represent the standard deviation to the mean of three biological replicates performed in triplicates on different days. **(D)** The two-dimensional response of the PJ23104 biosensor to glucose and *L*-lactate under aerobic and anaerobic conditions.

**Figure S3**

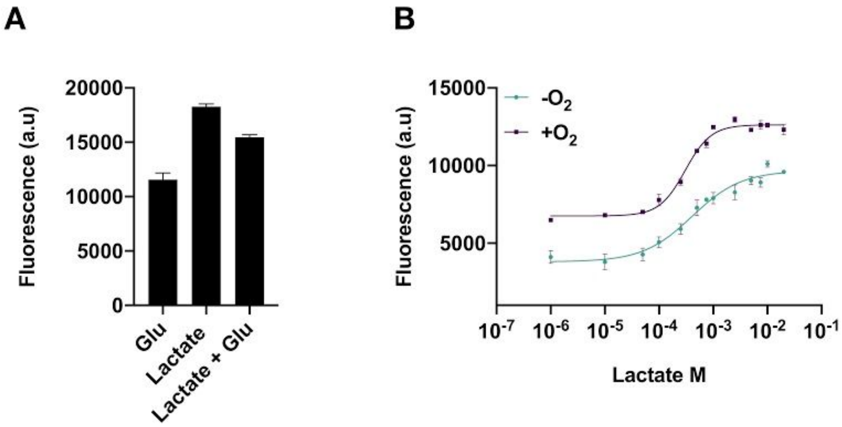


**Figure S3. Characterization of the first version of the synthetic L-lactate promoter showing high-leakiness (A)** Fluorescence (a.u.) response of the biosensor to 10 mM of L-lactate in the presence or not of 0.4% glucose. **(B)** The response function of the synthetic promoter lactate biosensor to L-lactate in presence of 0.4% of glucose in aerobic (+$O_2$) and (-$O_2$) anaerobic conditions.

**Figure S4**

**A**

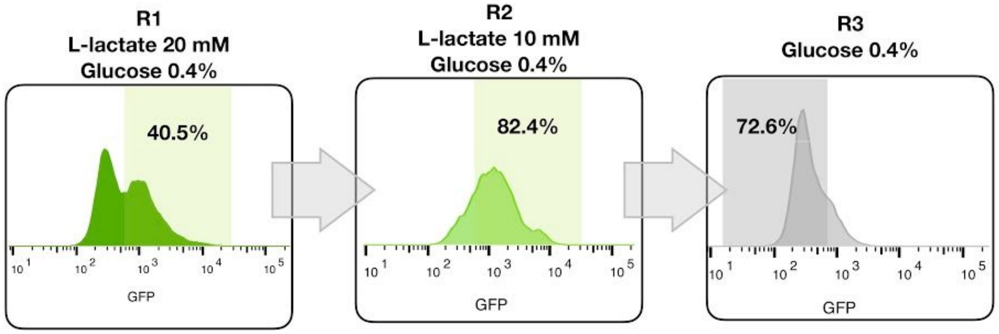


**B**

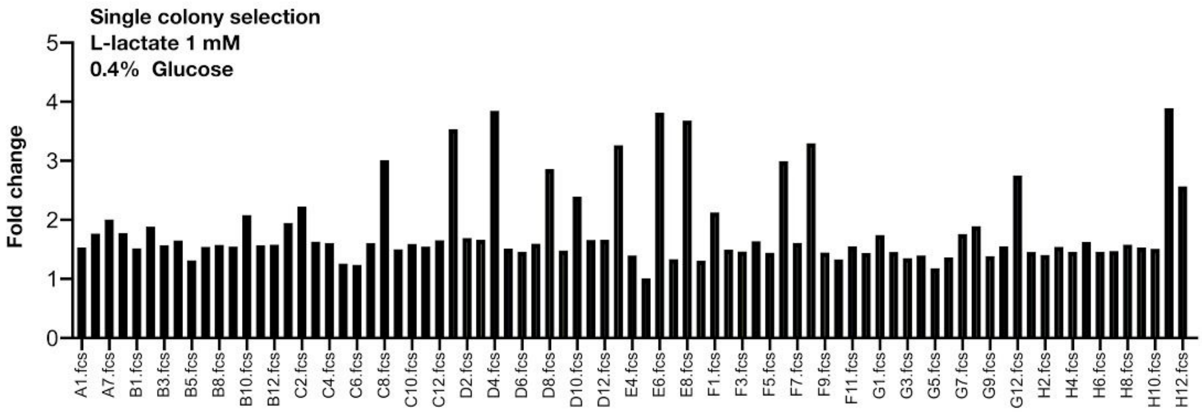


**Figure S4. Schematics of FACS enrichment and screening of the RBS library. (A)** The gates used for cell sorting are colored. For the first rounds of selection, the RBS library was induced with 20mM of lactate, the second with 10mM and in the third round, the cells were not induced to select those less leaky. **(B)** The final round was grown and plated with several dilutions on LB plates with kanamycin. 80 colonies were selected and induced for 4h with 1mM of lactate and glucose 04.% to determine their performance. Those with higher fold change were chosen and sequenced.

**Figure S5**

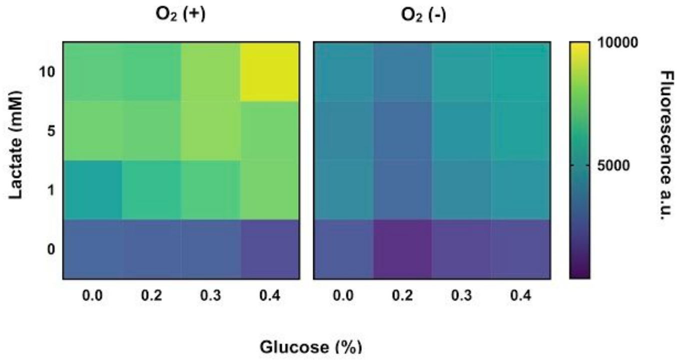


**Figure S5. Two-dimensional response of the ALPaGA L-lactate biosensor to increasing concentrations of lactate and glucose in aerobic (+$O_2$) and (-$O_2$) anaerobic conditions**.

**Figures S6**

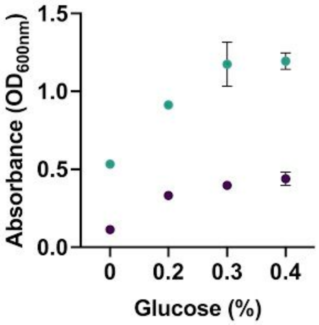


**Figure S6. Effect of glucose on the growth of the ALPaGA L-lactate biosensor.** Cells were induced for 16h with 10 mM of L-lactate in aerobic (green dots) or anaerobic (purple dots) conditions.
